# Acetylation of L-leucine switches its carrier from the L-amino acid transporter (LAT) to organic anion transporters (OAT)

**DOI:** 10.1101/2020.11.12.379701

**Authors:** Grant C. Churchill, Michael Strupp, Tatiana Bremova-Ertl, Mallory Factor, Marc C Patterson, Frances M. Platt, Antony Galione

**Author notes:** Corresponding author, (GCC).

## Abstract

N-acetyl-DL-leucine is an analogue of the alpha amino acid leucine with a chiral stereocenter. The active L-enantiomer of the racemate is currently under development for rare neurological disorders. Here we present evidence that a selective recognition of N-acetyl-L-leucine versus L-leucine by different uptake transporters significantly contributes to the therapeutic effects of N-acetyl-L-leucine. A previous study of the pharmacokinetics of racemic N-acetyl-DL-leucine and N-acetyl-L-leucine revealed D-L enantiomer competition and saturation kinetics, best explained by carrier-mediated uptake. The strategy we used was to first analyze the physicochemical properties associated with good oral bioavailable drugs and how these are alerted by N-acetylation by comparing N-acetyl-L-leucine with L-leucine. Using *in silico* computational chemistry we found that N-acetylation has a profound impact on certain physicochemical properties that can rationalize why N-acetyl-L-leucine is drug-like compared to L-leucine. Our calculations show that at physiological pH, L-leucine is a zwitterion, whereas N-acetyl-L-leucine is present as mainly an anion. Specifically, N-acetylation removes a charge from the nitrogen at physiological pH and N-acetyl-L-leucine is an anion that is then a substrate for the organic anion transporters. We examined N-acetyl-L-leucine uptake in human embryonic kidney cells overexpression candidate organic anion transporters (OAT) and pharmacological inhibitors. We found that N-acetyl-L-leucine is a translocated substrate for OAT1 and OAT3 with low affinity (Km ~10 mM). In contrast, L-leucine is known to be transported by the L-type Amino Acid Transporter (LAT) with high affinity (Km ~0.2 mM) and low capacity. The clinical consequence is that L-leucine uptake becomes saturated at 50-fold lower concentration than N-acetyl-L-leucine. These results demonstrate a mechanism of action that explains why N-acetyl-L-leucine is effective as a drug and L-leucine itself is not.

## INTRODUCTION

N-acetyl-DL-leucine has been used as an over-the-counter drug for the treatment of vertigo since 1957 (Tanganil^®^, Laboratoires Pierre Fabre; Neuzil et al., 2002). Currently, N-acetyl-leucine is being intensively studied by both academia and industry as a promising treatment for several disorders with unmet medical needs including cerebellar ataxia (Feil et al., 2017; Kalla and Strupp, 2019; Schniepp et al., 2016; Strupp et al., 2013), cognition and mobility in the elderly (Platt and Strupp, 2016), lysosomal storage disorders (Bremova et al., 2015; Cortina-Borja et al., 2018; Kaya et al., 2020a, 2020b), migraine (Strupp et al., 2019) and restless legs syndrome (Schoser et al., 2019). Three multinational clinical trials are ongoing with N-acetyl-L-leucine for the treatment of Niemann-Pick disease type C, GM2 Gangliosidosis, and Ataxia-Telangiectasia (Fields et al., 2020) (clinicaltrials.gov NCT03759639, NCT03759665, NCT03759678).

Given the promise of N-acetyl-L-leucine as a drug for treating many disease indications, there is intensifying interest in its mechanism of action. Although the pharmacodynamic mechanism of action of N-acetyl-leucine is not precisely known for any disease indication, for vertigo evidence implicates direct action on the central vestibular-related pathways and ocular motor networks (Kalla and Strupp, 2019; Vibert and Vidal, 2001), as also demonstrated by Positron Emission Tomography with [^18^F]fluorodeoxyglucose (Günther et al., 2015). At the molecular level, several mechanisms of action have been suggested including physicochemical partitioning into the phopspholipid bilayer to decrease its fluidity (Neuzil et al., 2002), direct action on glycine recptors and AMPA receptors (Neuzil et al., 2002), effects on branched chain aminotransferaes affecting glutamate neurotransmission (Kalla and Strupp, 2019), an increase in glucose metabolism (Kalla and Strupp, 2019). For the lysosomal storage disorder Niemann-Pick type C (NPC), N-acetyl-L-leucine mediates a shift in global cellular metabolism from glycolysis to oxidative phosphorylation for ATP production (Kaya et al., 2020), and in another lysosomal storage disorder Sandoff disease, it normalizes the aberant metabolism that results in oxidative stress (Kaya et al., 2020). All of the above mechanisms could contrribute to the observed effect of normalizing membrane potential and excitability leading to activation of the vestibulo-cerebellum and a deactivation of the posterolateral thalamus (Ferber-Viart et al., 2009; Kalla and Strupp, 2019; Vibert and Vidal, 2001). The complex and pleotropic pharmacodynamics of N-acetyl-L-leucine echo those of the actions of L-leucine inside cells (Pedroso et al., 2015). It is not clear if the effects of N-acetyl-L-leucine are due to the compound per se or whether it delivers L-leucine more effectively to tissues through an oral route of administration. In all the pharmacological studies, it was assumed that the applied compound was the active compound. This does not take into account the complexity of the interplay among transporters and metabolizing enzymes of N-acetyl-L-leucine. The best working hypothesis to reconcile the pharmacokinetic and pharmacodynamics data is that N-acetyl-L-leucine is functioning as via metabolic products (Churchill et al., 2020).

Therefore, in addition to the above pharmacodynamic effects of N-acetyl-L-leucine, pharmacokinetic effects may be playing a major contributing role to its mechanism of action and efficacy as a drug. Indeed, the combination of poor absorption and related poor pharmacokinetics is one of the main reasons for attrition in the drug development process (van De Waterbeemd et al., 2001). Moreover, our recent findings that the enantiomers of N-acetyl-leucine (N-acetyl-L-leucine and N-acetyl-D-leucine) show unexpected and large differences in pharmacokinetics (Churchill et al., 2020) supports the importance of pharmacokinetics in the mechanism of action of N-acetyl-L-leucine. Specifically, if enantiomers show pharmacokinetic differences, there must be biological sites of recognition that can discriminate between them such as enzymes and carrier-mediated uptake by transporters.

Transporters exist in the membrane of all cells and are required for the uptake of drugs that are not sufficiently hydrophobic to cross the membrane by passive diffusion (International Transporter Consortium et al., 2010; Keogh, 2012). As the major barrier to membrane crossing is the hydrophobic interior (Missner and Pohl, 2009), neutral molecules can pass by passive diffusion (Fig. 1E), whereas cations, anions and zwitterions (all alpha amino acids) can only cross with the aid or Solute Carriers (SLC) transporters (Fig. 1E). Approximately 450 transporter-like genes are expressed in humans and are categorized into two major superfamilies: the solute carrier (SLC) and ATP-binding cassette (ABC) transporters (Keogh, 2012). These superfamilies are categorized based on their gene sequence similarity, but exist in parallel with a historical classification system based on function and substrate specificity. For example, the organic anion transporters, where OAT1 is also known as SLC22A6 (VanWert et al., 2010).

**Figure 1.**
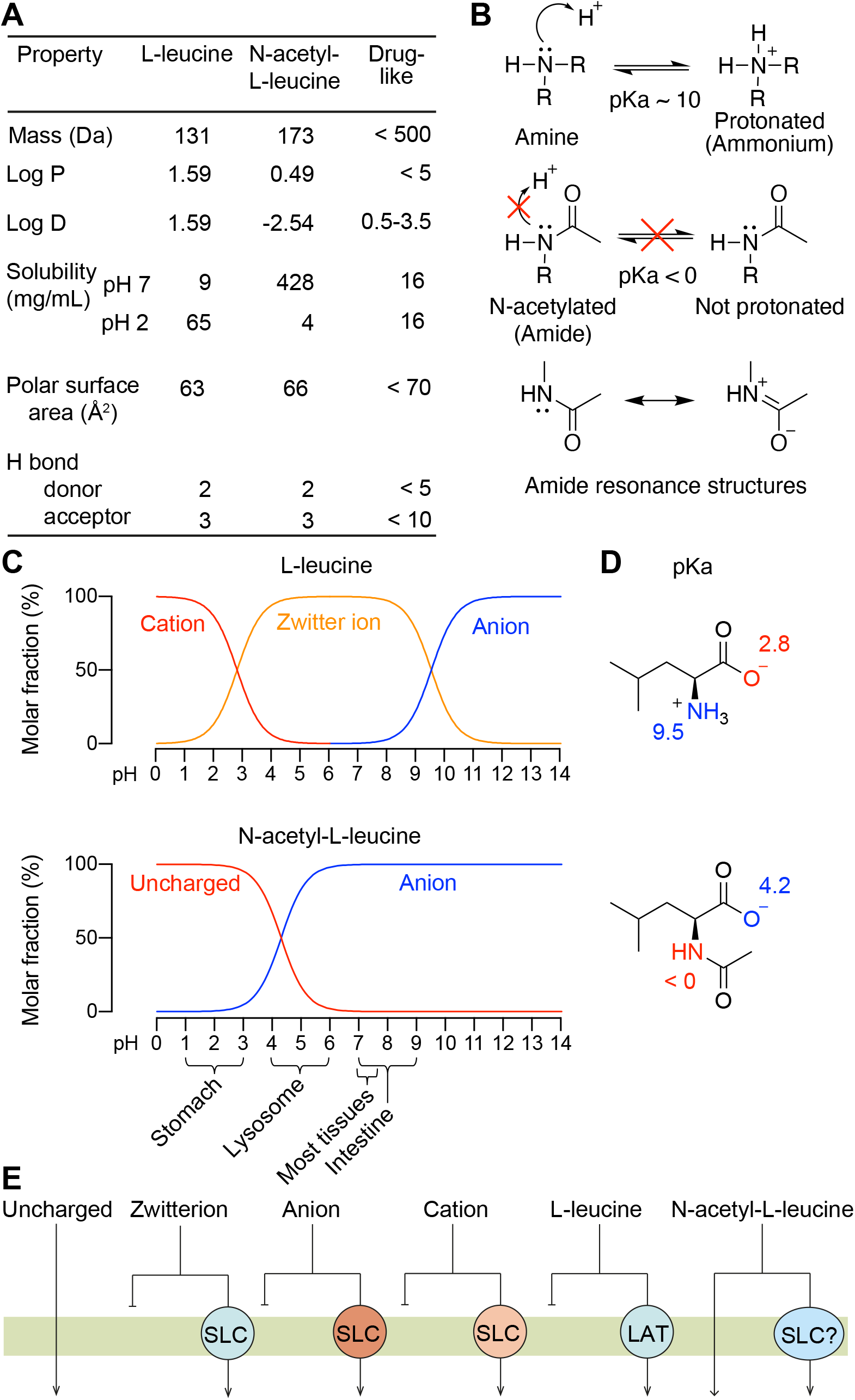
The effects of N-acetylation on the chemical properties and pharmacological consequences of the drug N-acetyl-L-leucine. (**A**) Comparison of the physicochemical properties of L-leucine and N-acetyl-L-leucine relating to oral bioavailability. (**B**) Lewis structures illustrating the effect of N-acetylation on the pK_a_ of nitrogen atom. Resonance delocalization of the lone pair electrons in the amide greatly decreases the basicity of the nitrogen relative to the amine, making the molecule neutral or charged, respectively. (**C**) Speciation curves for the protonation states of L-leucine and N-acetyl-L-leucine. The gross charge distribution of a molecule as a function of pH is calculated as well. The dominant species is indicated in several tissues relevant to drug absorption and distribution. (**D**) Chemical structures showing charge at pH 7 with the pKa of the amino and carboxylic acid groups labelled. (**E**) Mechanisms of absorption illustrated by crossing a membrane by passive diffusion or carrier-mediated uptake. In general, uncharged molecules can cross lipid bilayer membranes through simple passive diffusion, whereas charged molecules including zwitterions (a positive and negative charge within the same molecule), anions and cations cannot cross membranes without a transporter. Over 400 Solute Carrier (SLC) transporters are known with broad but overlapping selectivities for substrate. L-leucine as a obligate zwitterion at all biologically relevant pH values has an absolute requirement for its carrier, the high-affinity and low capacity L-amino Acid Transporter (LAT). In contrast, N-acetyl-L-leucine can exist as a neutral species and passively cross membranes at low pH, or as an anion recognized by other transporters. Physicochemical properties were calculated with Chemicalize provided by ChemAxon (https://chemicalize.com/app/calculation).

As transporters are often a rate-limiting step in drug absorption and distribution (International Transporter Consortium et al., 2010; Keogh, 2012), we sought to further investigate the uptake of N-acetyl-L-leucine by transporters. Our previous data indicates that uptake and distribution of N-acetyl-L-leucine involves transporters of low affinity and high capacity (Churchill et al., 2020), but the actual identity of such transporters are unknown. The L-amino Acid Transporter (LAT1/ SlC7A5) that transports L-leucine (Kanai et al., 1998; Krehbiel and Matthews, 2003) can be ruled out as it is high affinity and low capacity and does not bind or transport N-acetyl-L-leucine (Nagamori et al., 2016).

The objective of the present study was to explore uptake of N-acetyl-L-leucine. The strategy we used was to first analyze the physicochemical properties associated with good oral bioavailable drugs (Lipinski, 2000) and how these are alerted by N-acetylation by comparing N-acetyl-L-leucine with L-leucine. Using *in silico* computational chemistry we illustrate that N-acetylation has a profound impact on certain physicochemical properties that can rationalize why N-acetyl-L-leucine is drug-like and L-leucine is not. Our calculations show that at physiological pH, L-leucine is a zwitterion, whereas N-acetyl-L-leucine is present as mainly an anion. We then tested candidate organic anion transporters and found that N-acetyl-L-leucine is a translocated substrate for OAT1 and OAT3. These results provide a mechanistic explanation for why N-acetyl-L-leucine is an orally bioavailable drug and L-leucine is not. Specifically, N-acetylation shifs uptake of L-leucine from a high affinity and low capacity transporter (LAT) to low affinity and high capacity transporters that is about fifty-fold less saturable.

## MATERIALS AND METHODS

### General chemicals

High pressure liquid chromatography (HPLC) grade methanol and acetonitrile were obtained from Merck (Darmstadt, Germany), and formic acid, acetic acid and ammonium formate were obtained from BDH Laboratory Supplies (Poole, UK). N-Acetyl-L-Leucine was obtained from Laboratories Pierre Fabre and dissolved directly to incubation buffer at the highest incubation concentration on the day of incubations. All other chemicals were obtained from Sigma Aldrich (Helsinki, Finland) at the highest purity available. Water was in-house freshly prepared with a Direct-Q3 (Millipore Oy, Espoo, Finland) purification system and UP grade (ultra-pure, 18.2 MΩ).

### Human Solute Carrier (SLC) transporter-mediated cellular uptake

Human OAT1 (SLC22A6), OAT3 (SLC22A8) and OCT2 (SLC22A2) overexpressing HEK-293 cells and control cells without transfected transporter (Corning^®^ TransportoCellsTM) were plated to 24-well plates and cellular uptake of 10 to 100 μg/mL N-acetyl-L-leucine was measured in the absence and presence of transporter inhibitors. N-acetyl-L-leucine uptake into control cells was measured only without chemical inhibitors. Positive control substrates were incubated in parallel to demonstrate presence of active transport in each transporter transfected cell line.

DMEM (Gibco 4196, high glucose, without sodium pyruvate) supplemented with MEM nonessential amino acids and 10% fetal bovine serum. Cells were re-fed with fresh medium (supplemented with 2 mM sodium butyrate for OATP1B1 and OATP1B3 cells) after attachment (4-6 hours post-seeding). Cell were plated at a density of 4 × 10^5^/well in 24-well plates coated with poly-D-lysine. Transporter assays were conducted in 400 μL of HBSS supplemented with 10 mM Hepes, pH 7.4

The known probe substrates used as positive controls were 3 μM chlorothiazide for OAT1, 2 μM estrone-3-sulfate for OAT3, 30 μM metformin for OCT2, 100 μM diclofenac for OAT1, 100 μM diclofenac for OAT3 and 300 μM verapamil for OCT2. The known inhibitors used as controls for each transporter were 3 μM cyclosporine A for OATP1B1, 3 μM cyclosporine A for OATP1B3, 100 μM diclofenac for OAT1, 100 μM diclofenac for OAT3 and 300 μM verapamil for OCT2. All incubations were in triplicate and contained 1% DMSO. The duration of uptake was 5 min for OATP1B1, OAT1, OAT3 and OCT2 or 3 min for OATP1B3. Assays were performed at 37°C with no shaking. To terminate uptake the plate was placed on ice and cells washed twice with ice-cold transport buffer. To collect cells, they were detached with trypsin, and samples of cell suspension transferred into an equal volume of ice-cold acetonitrile. Samples were stored at −20°C until analysis.

In preparation for analysis, samples were centrifuged 20 min (4000 rpm) to separate the precipitated protein. Samples of supernatant were diluted 1:4 with acetonitrile (OCT2) or 1:4 with phosphate buffered saline (OATP1B1, OATP1B3, OAT1, OAT3). Samples diluted with phosphate buffered saline were used for analysis of N-acetyl-L-leucine (OCT2 samples split in two prior to dilution).

### Inhibition of Solute Carrier (SLC) transporters

Human OATP1B1 (SLCO1B1), OATP1B3 (SLCO1B3), OAT1 (SLC22A6), OAT3 (SLC22A8), OCT2 (SLC22A2) overexpressing HEK-293 cells (Corning^®^ TransportoCellsTM) were plated to 24-well plates and cellular uptake of probe substrate was measured in the absence and presence of N-acetyl-L-leucine at concentration ranging from 30 to 9000 μg/mL in conditions outlined below. Positive control inhibitors were incubated in parallel to estimate the passive cellular uptake of the probe substrate. Cells were seeded and cultured as described above for cellular uptake.

The probe substrates were 2 μM estradiol-17β-glucuronide for OATP1B1, 3 μM rosuvastatin for OATP1B3, 3 μM chlorothiazide for OAT1, 2 μM estrone-3-sulfate for OAT3 and 30 μM Metformin for OCT2. The known inhibitors used as controls for each transporter were 3 μM Cyclosporine A for OATP1B1, 3 μM Cyclosporine A for OATP1B3, 100 μM Diclofenac for OAT1, 100 μM Diclofenac for OAT3 and 300 μM verapamil for OCT2. Cells were preincubated for 0 and 30 min with N-acetyl-L-leucine for OATP1B1, OATP1B3 and for 0 min for OAT1, OAT3, OCT2. All incubations were in triplicate and contained 1% DMSO. The duration of uptake was 5 min for OATP1B1, OAT1, OAT3 and OCT2 or 3 min for OATP1B3. Assays were performed at 37°C with no shaking. To terminate uptake the plate was placed on ice and cells washed twice with ice-cold transport buffer. To collect cells, they were detached with trypsin, and samples of cell suspension transferred into an equal volume of ice-cold acetonitrile. Samples were stored at −20°C until analysis. Once thawed, samples were centrifuged 20 min (4000 rpm) to separate the precipitated protein. Samples of supernatant were diluted 1:4 with acetonitrile (OCT2) or 1:4 with phosphate buffered saline (OATP1B1, OATP1B3, OAT1, OAT3). Samples diluted with phosphate buffered saline were used for analysis of N-acetyl-L-leucine (OCT2 samples split in two prior to dilution).

### Liquid chromatography-mass spectrometry

N-acetyl-L-leucine, rosuvastatin, estrone-3-sulfate and chlorothiazide were separated and quantified using a Thermo Vanquish UPLC + Thermo Quantis triple quadrupole MS on a Waters Acquity HSS T3 (2.1 × 50 mm, 1.7 μm) column with guard filter. A sample of 4 μL was injected and compounds were eluted at 35°C with a flow of 0.65 mL/min using a gradient of solvent A = 0.1% formic acid and solvent B = acetonitrile as follows (Time, A%): 0.0, 95; 0.5, 95; 2.5, 40; 3.5, 5; 4.5, 95.

Metformin was separated and quantified using a Waters Acquity UPLC + Waters XEVO triple quadrupole MS on a Waters Acquity BEH HILIC (2.1 × 50 mm, 1.7 μm) column with guard filter. A sample of 4 μL was injected and compounds were eluted at 35°C with a flow of 0.5 mL/min using a gradient of solvent A = 10 mM ammonium formate and solvent B = acetonitrile as follows (Time, A%): 0.0, 5; 0.5, 5; 2.5, 50; 3.5, 50; 4.5, 5.

### Calculations

The IC_50_ value for the test item was determined by fitting the Hill equation in the following form:

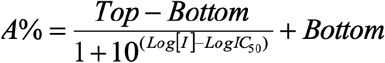

where *A*% is the percent activity remaining (the mean cellular uptake observed in the solvent control sample set to 100% and the mean cellular uptake observed in the presence of the positive control inhibitor set to 0%), *Top* and *Bottom* are the upper and lower plateau of A%. *[I]* is the inhibitor concentration and IC_50_ is the inhibitor concentration where the remaining activity is at the midpoint between the *Top* and *Bottom*. To obtain robust IC_50_ fit with four test concentrations the *Top* and the *Bottom* levels were constrained to 100% and 0%, respectively.

Enzyme kinetic data for N-acetyl-L-leucine uptake was analysed by fitting the Michaelis-Menten equation to the data. V_0_ = Vmax.[S]/Km + [S], where V_0_ is initial velocity, [S] is substrate concentration, Vmax is maximum velocity and Km is substrate concentration at half Vmax. All fitting was performed using GraphPad Prism 8.4 software (GraphPad Software Inc). No weighting scheme was applied.

### *In silico* chemical calculations

The physicochemical properties, pKa and the pH-dependent speciation curves for N-acetyl-L-leucine were performed with the online computational chemistry package Chemicalize (ChemAxon. https://chemicalize.com/app/calculation).

## RESULTS

### Analysis of the physicochemical properties associated with drug-like molecules

Several physicochemical parameters have been reported to correlate with the oral bioavailability of drugs (Camenisch et al., 1996; Gleeson et al., 2011; Leeson and Springthorpe, 2007; Lipinski, 2000; Waring, 2009; van de Waterbeemd et al., 1998). These physicochemical parameters relate to whether a small molecule can cross a membrane by passive diffusion, known in the field as being ‘drug-like’, or requires carrier-mediated transport (Fig. 1E; (van de Waterbeemd et al., 1998). Therefore, it is instructive to compare the amino acid L-leucine with the modified amino acid N-acetyl-L-leucine to help understand how N-acetylation converts a L-leuince into a drug. A comparison of N-acetyl-L-leucine with L-leucine reveals that both molecules possess similar physicochemical parameters, except for log P, log D and solubility (Fig. 1A). Log P is the octanol: water partition coefficient for the neutral form of a compound. In contrast, log D considers the ionization state of a molecule in aqueous (biological) solution resulting from basic groups gaining a proton and acidic groups losing a proton, and as such is better correlated with passive diffusion across membranes (Waring, 2009; van de Waterbeemd et al., 1998). For N-acetyl-L-leucine a log D of −2.54 (Fig. 1A) means that in a two phase octanol: water partition experiment for every molecule in octanol there would be 350 in water. As only the species in the octanol would be membrane permeant, N-acetyl-L-leucine have a low rate of passive diffusion across membranes (Gleeson et al., 2011; Lipinski, 2000; Missner and Pohl, 2009; Waring, 2009).

### Effect of N-acetylation on physicochemical and pharmacological properties

The physicochemical differences arising from N-acetylation can be understood from an analysis of the underlying chemistry. As expected for an amino acid with a non-ionizable side chain, L-leucine’s log P and log D are both 1.59 (Fig. 1A). In contrast, N-acetyl-L-leucine has a log P of 0.49 but a log D of log D of −2.54, making it 1000-fold less hydrophobic and 50-fold more soluble at the physiological pH of 7 compared to L-leucine (Fig. 1A). This is due to the effect of N-acetylation on the acid-base chemistry. In alpha amino acids the amino and a carboxylic acid group are bound to the same carbon, facilitating intramolecular interactions that affect both ionization state and hydrophobicity. At physiological pH, the amino acid exists as a zwitterion with a carboxylate and an ammonium (Fig. 1B). The charged groups interact through inductive effects of the electronegative nitrogen withdrawing charge as well as charge-charge coulombic electrostatic attraction and stabilization. As the charges interact with one another they are not available for interacting with water molecules, making the compound more hydrophobic and less soluble than expected (Fig. 1A). The pK_a_ of a carboxylic acid in an amino acid is 2-3, making it 100-fold more acidic than carboxylic acid functional groups bound to an alkane chain (Fig. 1D). Upon N-acetylation the carboxylic acid’s pK_a_ shifts from 2.8 to 4.2 (Fig. 1D).

The other major effect of N-acetylation is on the acid-base properties of the nitrogen. In an amino acid, the lone pair of electrons is basic and protonated with a pK_a_ of about 10 (Fig 1B and D). N-acetylation forms an amide bond, which is much less basic as the pK_a_ shifts from 10 to less than 1 (Fig. 1D). This can be best described through molecular orbital theory (Hall, 1991) but more intuitively through Lewis structures (Lewis, 1916) based on valance bond theory and resonance (Pauling, 1977) in which the lone pair electrons participate in binding and are partially delocalized to the adjacent carbon in the N-C bond and the carbonyl oxygen making it a partial double bond (Fig. 1B), resulting in less electron density being available for binding to protons (Fig. 1B). The differing pK_a_ of the nitrogen and oxygen have profound effects on the ionization state of the two molecules. L-leucine has two pK_a_ values and can be present as a cation, zwitterion and anion, but is predominately a zwitterion at physiological pH 7 (Fig. 1C). In contrast, N-acetyl-L-leucine has a single pK_a_ and can exist as a neutral molecule, but is predominately an anion at physiological pH 7 (Fig. 1C). The pharmacological consequence is that as an anion, N-acetyl-L-leucine requires a transporter for uptake.

### N-acetyl-L-leucine inhibits the solute carrier (SLC) transporters OAT1, OAT3 and OCT2

Transport of a small molecule can be modelled and analysed as a two-step process in which a molecule first binds (affinity) and then is translocated across the membrane. That is, a molecule can be recognized only (inhibitor) or recognized and transported (substrate), and, as dictated by mass action competition for a single binding site, a substrate would be an inhibitor of another substrate. To determine whether N-acetyl-L-leucine was interacting with Solute Carrier (SLC) transporters, the ability of N-acetyl-L-leucine to act as an inhibitor was evaluated. These inhibition experiments used a known substrate (probe) of a given transporter to measure the effect of N-acetyl-L-leucine at various concentrations (Fig. 2A).

**Figure 2.**
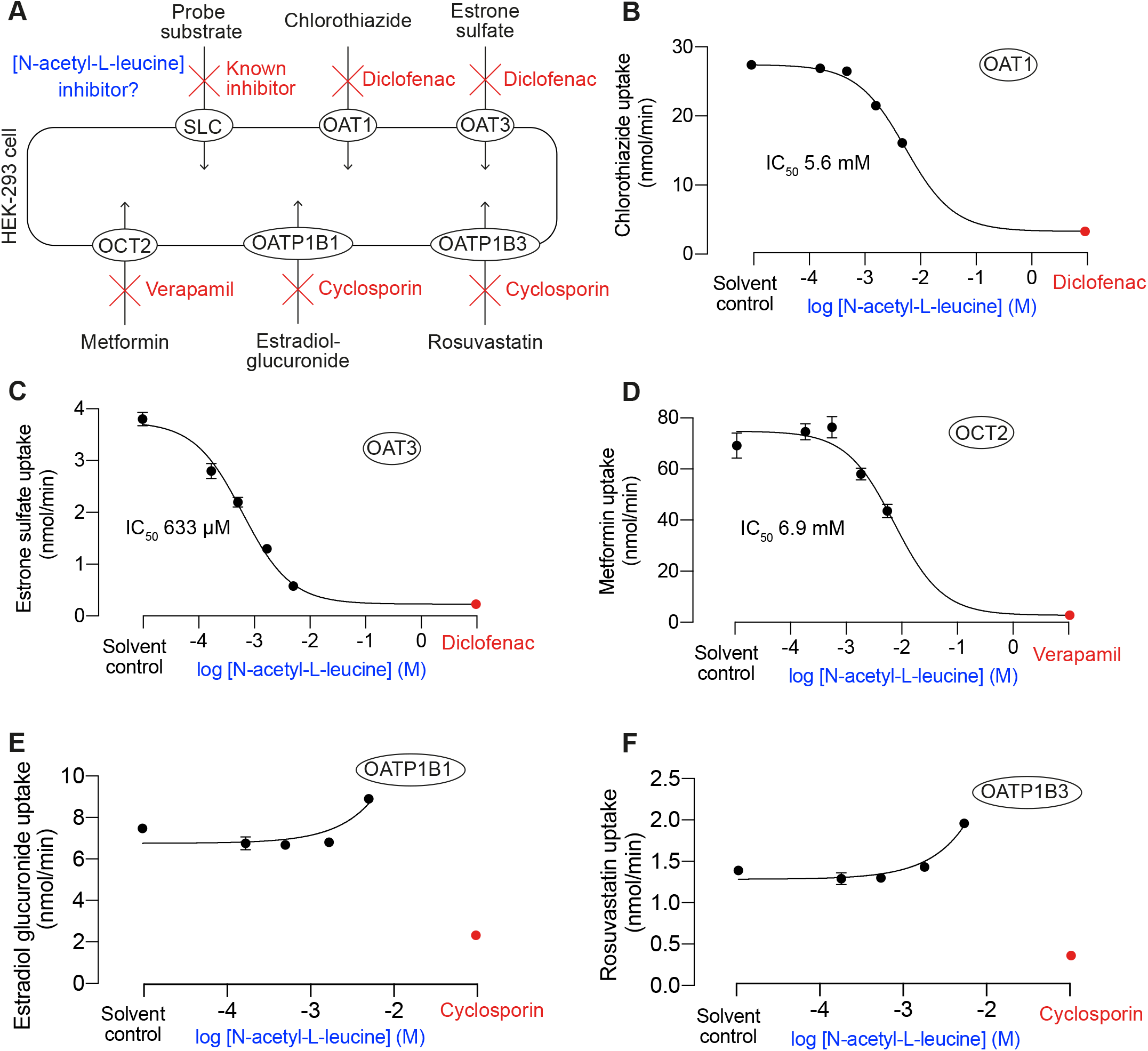
Evaluation of inhibition of Solute Carrier (SLC) transporters by L-leucine and N--acetyl-L-leucine in vitro. (**A**) Schematic showing how transporter activity was determined by uptake of a known probe substrate into cells over expressing a human SLC transporter in HEK-293 cells. Inhibition of active transport was observed as decreased cellular uptake of the probe substrate in the presence of inhibitor. The effect of N-acetyl-L-leucine concentration on the transport activity of (**B**) OAT1, (**C**) OAT3, (**D**) OCT2, (**E**) OATP1B1 and (**F**) OATP1B3. Data were fit to the Hill equation using the solvent control (DMSO 1%) to define the top and the positive control inhibitor to define the bottom. Symbols represent the mean ± SEM, n = 3. When the error bars are smaller than the symbol, they are not visible.

In these inhibition experiments N-acetyl-L-leucine was tested at 30, 90, 300, 900, 3000 and 9000 μg/mL corresponding to 0.173, 0.520, 1.73, 5.20, 17.34 and 52.0. During the incubations with N-acetyl-L-leucine present at the two highest test concentrations, 17 and 52 mM (3000 and 9000 μg/mL), the cells appeared to detach from the wells during followed by tight binding to the wells after washing (based on subjective visual evaluation by naked eye). Furthermore, apparent probe substrate uptake results showed erratic apparent concentration dependence at these concentrations and were thus considered not reliable as a measure of specific transporter interaction. Consequently, transporter inhibition was evaluated based on results from incubations at 4 concentrations of N-Acetyl-L-Leucine ranging from 0.173 to 5.2 mM (30 to 900 μg/mL).

N-acetyl-L-leucine inhibited the activity of the organic anion transporters (OAT) OAT1 (Fig. 2B) and OAT3(Fig. 2C) with IC_50_ values (a proxy of affinity) of 5.6 mM and 0. 6 mM, respectively. N-acetyl-L-leucine also inhibited the activity of the organic cation transporter (OCT) OCT with IC_50_ of 6.9 mM (Fig. 2D). N-Acetyl-L-Leucine at concentrations from 0.734 to 1.73 mM did not affect activity of the organic anion transporting polypeptides (OATP) OATP1B1 (Fig. 2E) and OATP1B3 (Fig. 2F). Although N-acetyl-L-leucine appeared to promote the activity of OATP1B1 (Fig. 2E) and OATP1B3 at 5.2 mM (Fig. 2F) based on increased apparent cellular uptake of the OATP1B1 and OATP1B3 probe substrates. However, involvement of a non-specific mechanism, instead of direct OATP transporter activation, at high N-Acetyl-L-Leucine concentrations cannot be ruled out.

Apparent probe substrate cellular uptake rate in the presence of positive control inhibitors ranged from 0.23 to 2.73 pmol/min depending on tested cell lines whereas cellular uptake rate in the solvent control incubations was 5–25 fold faster than with positive control inhibitors. This confirms the activity of the overexpressed transporters in the assays and capability to detect transporter inhibition. The solvent control and positive control inhibitors were also used in the curve fitting to the Hill equation as recommended as the most robust method when the test substance (N-acetyl-L-leucine in our case) could not be used at a sufficiently high concentration to provide complete inhibition (Weimer et al., 2012).

In summary, N-acetyl-L-leucine inhibits OAT1, OAT3 and OCT2 in a concentrationdependent manner.

### N-acetyl-L-leucine is a substrate for the solute carrier (SLC) transporters OAT1 and OAT3

Having established that N-acetyl-L-leucine inhibits the transport of known specific substrates of three of the SLC transporters tested, we investigated whether N-acetyl-L-leucine is also a substrate, that is, does N-acetyl-L-leucine not only bind (has affinity) but it is also translocated across the plasma membrane. The experimental design for these uptake experiments is shown schematically in Fig. 3A. Cellular uptake of N-Acetyl-L-Leucine into single human transporter transfected cells was measured at two concentrations (10 and 100 μg/mL) in the absence and presence of chemical transporter inhibitors, and into control cells without transfected transporter in the absence of chemical inhibitors. Transporter mediated cellular uptake of the study compounds is observed as difference in the cellular uptake to transporter overexpressing cells and vector transfected control cells and as inhibition of cellular uptake in transporter overexpressing cells by the control inhibitors. Positive control substrates at single concentration were assayed in parallel.

**Figure 3.**
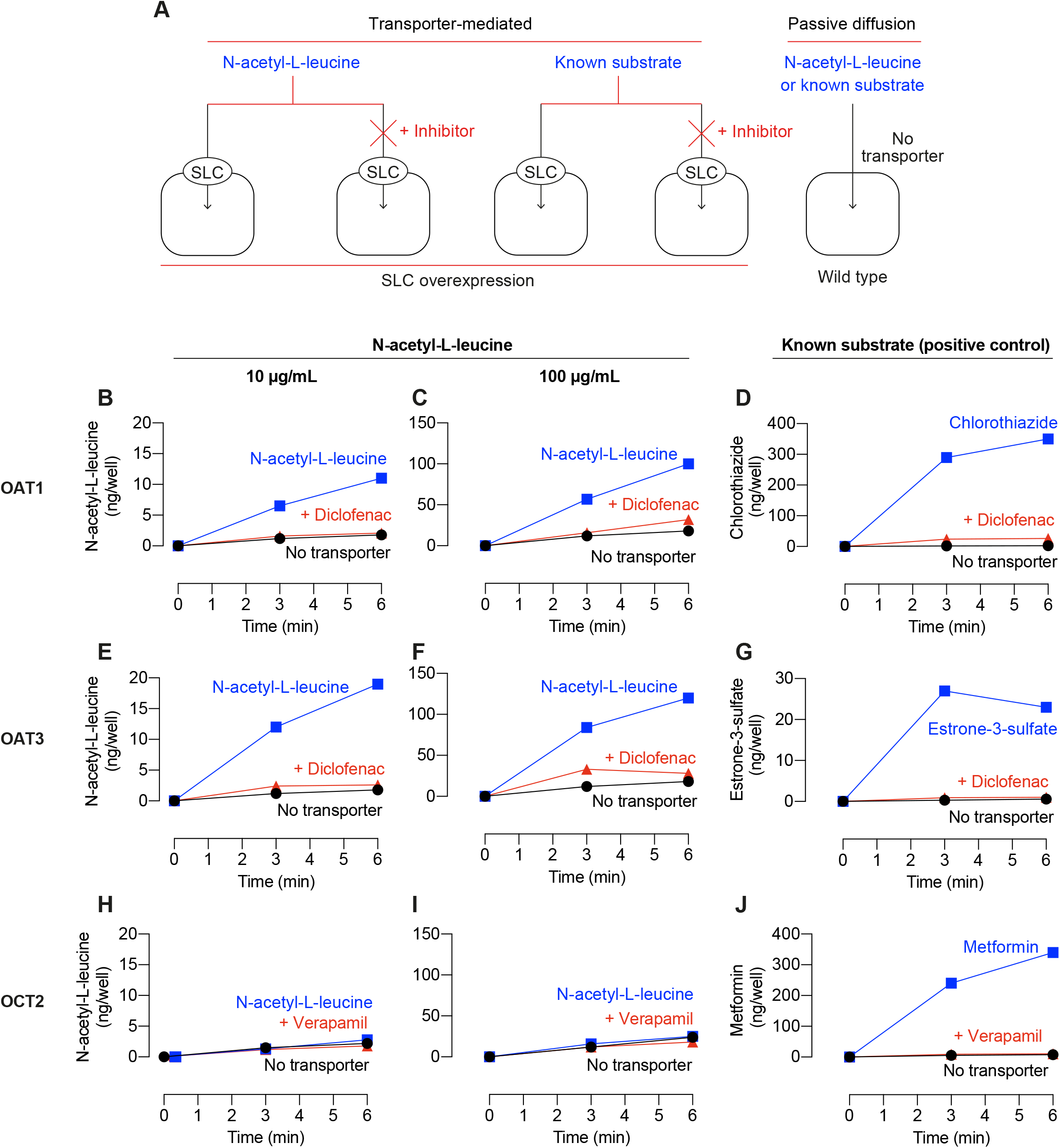
Time course of Solute Carrier (SLC) transporter-mediated uptake of N-acetyl-L-leucine into cells. (**A**) Schematic showing the experimental design. N-acetyl-L-leucine uptake was determined in HEK-293 cells overexpressing a single human SLC transporter protein in the absence and presence of known chemical transporter inhibitors, Wild type HEK-293 cells were used as controls (labelled No transporter). (**B-J**) Plots N-acetyl-L-leucine or known substrate uptake over time by cells expressing OAT1 (SLC22A6), OAT3 (SLC22A8) or OCT2 (SLC22A2). N-acetyl-L-leucine was present at 10 μg/mL or 100 μg/mL as indicated. A known substrate of each transporter was used as a positive control. Details of each experiment are shown as labelled on each panel. Symbols represent the mean ± SEM, n = 3. When the error bars are smaller than the symbol, they are not visible.

When present at 10 μg/mL (0.0578 mM), N-acetyl-L-leucine uptake in cells expressing the transporter OAT1 was about 5-fold higher at both 3 and 6 min compared both to cells not expressing the transporter and in the presence of the OAT1 inhibitor diclofenac (Fig. 3B). Similarly, when present at 100 μg/mL (0.578 mM), N-acetyl-L-leucine uptake was also about 5-fold higher at both time points compared to the controls (Fig. 3C). The 10-fold increase in N-acetyl-L-leucine concentration resulted in 10-fold more uptake (Fig. 2B compared to Fig. 2C), indicating that the transporters were not significantly saturating at 0.578 mM. The known positive control probe substrate chlorothiazide was taken up and inhibited by diclofenac (Fig. 3D), demonstrating the cellular uptake system was functioning properly. In summary, these data demonstrate that N-acetyl-L-leucine is a substrate for the OAT1 transporter.

When present at 10 μg/mL (0.0578 mM), N-acetyl-L-leucine uptake in cells expressing the transporter OAT3 was about 10-fold higher at both 3 and 6 min compared both to cells not expressing the transporter and in the presence of the OAT3 inhibitor diclofenac (Fig. 3E). Similarly, when present at 100 μg/mL (0.578 mM), N-acetyl-L-leucine uptake was about 7-fold higher at both time points compared to the controls (Fig. 3F). The 10-fold increase in N-acetyl-L-leucine concentration resulted in 7-fold more uptake (Fig. 2E compared to Fig. 2F), indicating that this transporter was beginning to saturate at 0.578 mM. The known positive control probe substrate estrone-3-sulfate was taken up and inhibited by diclofenac (Fig. 3G), demonstrating the cellular uptake system was functioning properly. In summary, these data demonstrate that N-acetyl-L-leucine is a substrate for the OAT3 transporter.

In cells expressing the transporter OCT2, N-acetyl-L-leucine was not taken up to a greater extent than in cells not expressing the transporter or in the presence of the inhibitor verapamil when present at either 10 μg/mL (Fig. 3H) or 100 μg/mL (Fig. 3I). The known positive control probe substrate estrone-3-sulfate was taken up and inhibited by diclofenac (Fig. 3J), demonstrating the cellular uptake system was functioning properly. In summary, these data demonstrate that although N-acetyl-L-leucine inhibits OAT3 (Fig. 2 D), it is not a substrate for the OAT3 transporter.

In all the N-acetyl-L-leucine uptake experiments (Fig. 3), a small amount of uptake was present in both the cellular negative control not expressing a transporter and in the positive control with a known transporter chemical inhibitor. As this uptake was not dependent on a transporter, it demonstrates N-acetyl-L-leucine uptake by passive diffusion.

To determine the kinetic parameters for N-acetyl-L-leucine uptake, the dependence of rate of uptake on concentration was investigated. In these kinetic experiments, N-acetyl-L-leucine was tested at 30, 90, 300, 900, 3000 and 9000 μg/mL corresponding to 0.173, 0.520, 1.73, 5.20, 17.34 and 52.0 mM. However, as mentioned above, the two highest concentrations were excluded due to suspected non-specific and/or toxic effects. Both the transporter OAT1 and OAT3 showed saturation kinetics and were well fit by the Michaelis-Menten equation with a K_m_ of 11 and 8 mM and V_max_ of 93 and 75 ng/min/well, respectively (Fig. 4). In summary, the kinetic experiments demonstrate transporter-mediated uptake of N-acetyl-L-leucine.

**Figure 4.**
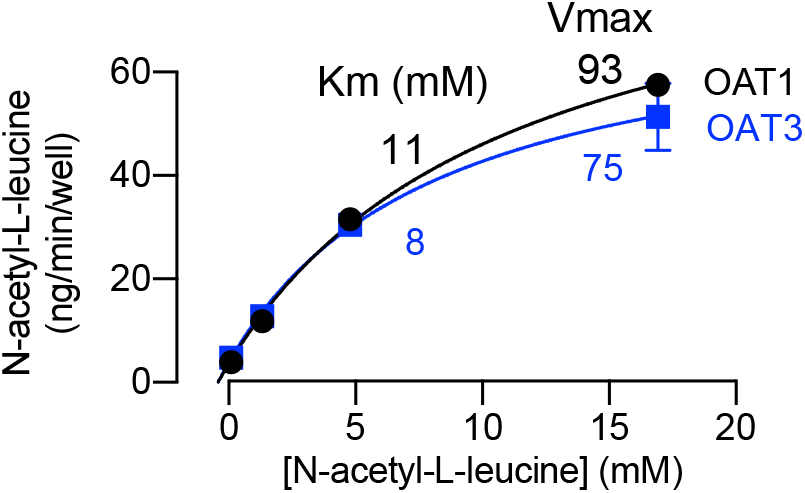
Concentration-dependent kinetics of N-acetyl-L-leucine by solute carrier (SLC) transporters. Data were obtained using HEK-293 cells overexpressing human organic anion transporter (OAT) OAT1 (SLC22A6), OAT3 (SLC22A8) or OCT2 (SLC22A2) incubated with the indicated concentration of N-acetyl-L-leucine for 5 min. Data were fit to the Michaelis-Menten equation to calculate the K_m_ and V_max_. Symbols represent the mean ± SEM, n = 3. When the error bars are smaller than the symbol, they are not visible.

## DISCUSSION

### General overview

As part of our continuing investigations into the drug N-acetyl-L-leucine, we explored its pharmacokinetics in regard to carrier-mediated uptake by transporters. Our previous work demonstrated differential uptake and mutual interference between the enantiomers of N-acetyl-leucine (Churchill et al., 2020). Enantiomeric differences only arise if there are biological recognition sites that can distinguish molecular chirality, meaning transporters and enzymes (Ariëns, 1986). In our current paper, we analysed the physicochemical and acidbase and ionization states of N-acetyl-L-leucine. We compared these to L-leucine, to gain insight into the effect of N-acetylation uptake. We demonstrated that at physiological pH N-acetyl-L-leucine exists as an anion, which would require carrier-mediated uptake across membranes. We then tested whether N-acetyl-L-leucine was recognized and a substrate for candidate organic anion transporters (OAT). N-acetyl-L-leucine is a transportable substrate of OAT1 and OAT3 with a Km of ~10 mM. This uptake mechanism and low affinity means that N-acetyl-L-leucine will not saturate its transporter and continue to be effective at concentrations about 100-fold over those for the unmodified amino acid L-leucine. The pharmacokinetics of transporter uptake by a low affinity process translates into levels of bioavailability of N-acetyl-L-leucine that are not possible with L-leucine, as its transporter saturates at concentrations of about 100-fold lower. The pharmacokinetics of N-acetyl-L-leucine explain why it is a drug, while the essential amino acid L-Leucine is not. This distinct transport mechanism provides a mechanistic explanation for the pharmacological and clinical relevance of N-acetyl-L-leucine over L-leucine itself.

### N-acetyl-L-leucine is a transport substrate of the organic anion transporters

Our determination that the major ionized form of N-acetyl-L-leucine at physiological pH (around 7) was an anion led us to test whether it was a substrate for organic anion transporters. We found that N-acetyl-L-leucine was a substrate of OAT1 and OAT3. OATs are expressed in the brain and kidney and is selective for small and hydrophilic molecules. OATs are organic anion exchangers that use an endogenous dicarboxylic acid from inside the cell such as glutamate or ketoglutarate. OATs have well-characterized structure-activity relationships (International Transporter Consortium et al., 2010; Kaler et al., 2007; Truong et al., 2008) and would be predicted to treat N-acetyl-L-leucine as a substrate. The mouse isoform, mOat1, prefers high polar surface areas (e.g. phosphate groups), whereas mOat3 prefers hydrogen bond acceptors (e.g. amines, ketones) and low rotatable bond numbers. OATs are well characterized and involved mostly in drug and nutrient distribution rather than uptake from the gastrointestinal tract (VanWert et al., 2010).

### Implications of transporter kinetics with high or low affinity

In general, the affinities of transporters the are considered to be of high affinity when K_m_ < 0.5 mM, medium affinity when K_m_ 0.5–5 mM and low affinity when K_m_ 5-15 mM (Brandsch et al., 2008). In general, there is often an inverse relationship and trade-off between affinity and capacity. In regard to biochemical processes, be they receptor binding of ligand, enzymes binding substrate or transporters binding substrate, a biophysical constraint is imposed by the theoretical maximum number of molecular collisions in a bimolecular reaction (k_on_) being diffusion limited at 10^8^ M^−1^ s^−1^ and plugged into the kinetic definition of affinity where K_d_ = k_off_/k_on_. Therefore, a high affinity process cannot be a fast or high-capacity process, whereas a low affinity process can be high or low capacity depending on efficiency of enzymatic reaction, transport or conformational change to an activated receptor. The biochemical and evolutionary advantages and disadvantages of high affinity/low capacity and low capacity/high affinity have been the subject of intense theoretical and experimental studies, with one view being that they favour scarce or plentiful nutrients, respectively (Levy et al., 2011). Far less attention has been given to drug transport, but the consequences can be profound, as we describe below.

Our current findings are consistent with the hypothesis that N-acetyl-L-leucine is a prodrug and is so effective because it can deliver L-leucine more effectively as it is handled by transporters with low affinities and high capacities (Churchill et al., 2020). L-leucine itself has potent physiological and biochemical effects but only when present *inside* cells (Pedroso et al., 2015). If L-leucine is the active compound, then transport by different transporters provides a satisfying mechanistic explanation for the efficacy of N-acetyl-L-leucine and the limitations of L-leucine.

As with all alpha amino acids, L-leucine is a zwitterion requires a carrier to cross membranes. This process is well characterized and uses the L-type Amino Acid Transporter (LAT1/SlC7A5), which is responsible for the intestinal uptake of L-leucine (Kanai et al., 1998; Krehbiel and Matthews, 2003). The LAT transporter has broad substrate promiscuity in that it transports D-leucine and is responsible for transporting several large neutral amino acids in addition to L-leucine including L-tryptophan, L-isoleucine and L-phenylalanine into cells (Kanai, 1998), and also transports amino acid-derived drugs such as levodopa, gabapentin, melphalan and baclofen (International Transporter Consortium et al., 2010; Krehbiel and Matthews, 2003). However, N-acetyl-L leucine is not an inhibitor or substrate (Nagamori et al., 2016). LAT is a high affinity and low capacity transporter (Kanai et al., 1998; Pedroso et al., 2015) with a K_m_ for L-leucine of 0.2 mM and D-leucine with a Km of 7.2 mM (Böhmer et al., 2005; Jørgensen et al., 1990), suggesting it is best suited for scavenging scarce nutrients (Levy et al., 2011). In summary, L-leucine is transported by a high affinity and low-capacity transporter which limits the maximum rate of uptake from the gastrointestinal tract and by tissues when the transporter is saturated with substrate.

That N-acetyl-L-leucine delivers more drug or prodrug to tissues than possible with L-leucine can be illustrated by a numerical example comparing K_m_ and compound concentrations. Taking stomach volume of a fasted individual as the accepted standard of 0.25 L (Pade and Stavchansky, 1998) a 4-g dose would result in 122 mM L-leucine and 92 mM N-acetyl-L-leucine. For L-leucine the LAT transporter would be 50% saturated at 0.2 mM (its Km), 90% saturated at 2 mM and 99% saturated at 20 mM, corresponding to oral ingestion of 0.000656 g, 0.00656 g and 0.656 g, respectively. In contrast, N-acetyl-L-leucine, if transported by OAT with a K_m_ of ~10 mM would it be 50% saturated at 10 mM (its K_m_), 90% saturated at 100 mM and 99% saturated at 1000 mM, corresponding to an oral ingestion of 0.43 g, 4.3 g and 43 g, respectively. Expressed another way, uptake of N-acetyl-L-leucine would continue at doses 66-fold higher than L-leucine. Therefore, even heroic concentrations of L-leucine outside cells would be limited in uptake by saturation of the high affinity LAT. This kinetic analysis holds for any low K_m_ transporter as observed for the absorption of N-acetyl-L-leucine which does not show saturation with a high dose (Churchill et al., 2020). This situation exists for uptake across any biological membrane, meaning the affinity and capacity of a transporter would limit oral absorption and tissue uptake in the whole organism. Moreover, even in cultured cells, the presence and level of transporter would affect the results and interpretation of experiments.

### Passive membrane permeability of N-acetyl-L-leucine

The analysis of the physicochemical properties of N-acetyl-L-leucine not only guided us to show that it is a translocated substrate for the organic anion transporters, but also reveals a contribution from passive diffusion across membranes. Uptake by passive diffusion is determined by physicochemical properties, primarily hydrophobicity (Camenisch et al., 1996; Missner and Pohl, 2009), and for ionisable molecules, only the neutral form is soluble in the membrane and is pH dependent. This is known as the pH partition hypothesis (Camenisch et al., 1996). The shift of the carboxylic acid pK_a_ to 4.2 (Fig. 1D) with N-acetylation results in a dramatically shifted speciation curve at various pH values compared to L-leucine (Fig. 1C). The pharmacologically relevant conclusion from analysis of the physicochemical properties of N-acetyl-L-leucine is that L-leucine that exists as a zwitterion and N-acetyl-L-leucine exists as an anion (Fig. 1D). This is pH dependent but holds for pH 7, for which most cells and body fluids exist, except for stomach contents (pH 1-3), lysosomes (pH 4-6) and the small intestine (pH 7-9) (Fig. 1C). Importantly, a neutral species of N-acetyl-L-leucine is present, the proportion of which depends on pH (Fig. 1C).

For N-acetyl-L-leucine, in the stomach (pH 1-3) the neutral species will predominate and be passively membrane permeable. However, the surface area for absorption in the stomach is far less (~ 1 m^2^) than that is the small intestine (~200 m^2^). At pH 7, 0.16% of N-acetyl-L-leucine would be uncharged and membrane permeant. In the plasma, this would correspond to 0.1 μM. In the small intestine, the total N-acetyl-L-leucine concentration would be 92 mM of which 15 mM would be membrane permeant. Comparing this to amino acids, which are present in the 50-100 μM range, this represents a very high cell permeable concentration of about 100-fold over endogenous amino acid concentrations (including L-leucine).

For the purposes of comparison, the passive permeability of amino acids such as L-leucine is similar to monovalent cations such as sodium and potassium with a permeability coefficient of approximately 10^−13^ cm.s^−1^ compared to that for water of 10^−2^ (Chakrabarti, 1994), as they would have to translocate in their neutral form, which is only 10^−7^ of the zwitterionic charged form at pH 7. Eliminating one of the two charges eliminates one of the charges present on an amino acid at physiological pH, which can increase transport rates up to 10^10^ fold (Chakrabarti, 1994).

In summary, passive diffusion across membranes likely contributes to the uptake and distribution of N-acetyl-L-leucine. As a small anion, the passive diffusion across membranes of N-acetyl-L-leucine would be expected to be very similar to that of aspirin with a pK_a_ of 3.5 and its pharmacokinetics are thought to be entirely due to passive diffusion of the neutral species across membranes (Levy, 1980).

### Conclusions and clinical implications

We have demonstrated carrier-mediated transport of N-acetyl-L-leucine by the organic anion transporters OAT1 and OAT3. As the kinetic constants for these transporters for N-acetyl-L-leucine are low affinity and high capacity, transporter uptake can provide a mechanistic explanation for why N-acetyl-L-leucine is effective as an oral therapeutic compared to L-leucine itself.

## Notes

### Competing Interest Statement

GCC is a cofounder, shareholder and consultant to IntraBio. MS is a shareholder to IntraBio, and consultant for Abbott, Actelion, AurisMedical, Heel, IntraBio and Sensorion; he has received speaker's honoraria from Abbott, Actelion, Auris Medical, Biogen, Eisai, Grunenthal, GSK, Henning Pharma, Interacoustics, Johnson & Johnson, MSD, Otometrics, Pierre-Fabre, TEVA, UCB. TBE received honoraria for lecturing from Actelion and Sanofi Genzyme. MF is a co-founder, shareholder, and Chairman of IntraBio. MCP is a shareholder of IntraBio, and has received consulting fees, honoraria and research grants from Actelion Pharmaceuticals Ltd. and Biomarin. FMP is a cofounder, shareholder, and consultant to IntraBio and consultant to Actelion and Orphazyme. AG is a cofounder, shareholder and consultant to IntraBio. IntraBio owns patents EP3359146 and EP3416631 (related to treatment of lysosomal storage disorders and neurodegenerative diseases with acetyl-Leucine and its analogues). IntraBio has pending patent applications EP19174007.5, EP3482754, PCT/US2018/056420, PCT/US2018/018420, PCT/IB2018/054676, PCT/IB2019/051214, PCT/IB2017/054928, PCT/GB2017/051090, PCT/IB2017/054929, USPTO 62/812,987, USPTO 62/842,296, USPTO 62/888,894, USPTO 62/895,144, USPTO 62/868,383, USPTO 62/931,003, USPTO 62/960,637, and PCT/IB2019/060525 relating to treatment of lysosomal storage disorders, neurodegenerative diseases and neurodegeneration with acetyl-leucine and its analogues.

## REFERENCES

Ariëns, E.J. (1986). Chirality in bioactive agents and its pitfalls. Trends Pharmacol. Sci. 7, 200–205.

Böhmer, C., Bröer, A., Munzinger, M., Kowalczuk, S., Rasko, J.E.J., Lang, F., and Bröer, S. (2005). Characterization of mouse amino acid transporter B0AT1 (slc6a19). Biochem. J. 389, 745–751.

Borgström, L., Kågedal, B., and Paulsen, O. (1986). Pharmacokinetics of N-acetylcysteine in man. Eur. J. Clin. Pharmacol. 31, 217–222.

Bowden, N.A., Sanders, J.P.M., and Bruins, M.E. (2018). Solubility of the proteinogenic α-amino acids in water, ethanol, and ethanol-water mixtures. J. Chem. Eng. Data 63, 488–497.

Brandsch, M., Knütter, I., and Bosse-Doenecke, E. (2008). Pharmaceutical and pharmacological importance of peptide transporters. J. Pharm. Pharmacol. 60, 543–585.

Bremova, T., Malinová, V., Amraoui, Y., Mengel, E., Reinke, J., Kolníková, M., and Strupp, M. (2015). Acetyl-dl-leucine in Niemann-Pick type C: A case series. Neurology 85, 1368–1375.

Camenisch, G., Folkers, G., and van de Waterbeemd, H. (1996). Review of theoretical passive drug absorption models: Historical background, recent developments and limitations. Pharm. Acta Helv. 71, 309–327.

Chakrabarti, A.C. (1994). Permeability of membranes to amino acids and modified amino acids: Mechanisms involved in translocation. Amino Acids 6, 213–229.

Churchill, G.C., Strupp, M., Galione, A., and Platt, F.M. (2020). Unexpected differences in the pharmacokinetics of N-acetyl-DL-leucine enantiomers after oral dosing and their clinical relevance. PloS One 15, e0229585.

Cortina-Borja, M., Te Vruchte, D., Mengel, E., Amraoui, Y., Imrie, J., Jones, S.A., I Dali, C., Fineran, P., Kirkegaard, T., Runz, H., et al. (2018). Annual severity increment score as a tool for stratifying patients with Niemann-Pick disease type C and for recruitment to clinical trials. Orphanet J. Rare Dis. 13, 143.

van De Waterbeemd, H., Smith, D.A., Beaumont, K., and Walker, D.K. (2001). Property-based design: optimization of drug absorption and pharmacokinetics. J. Med. Chem. 44, 1313–1333.

Feil, K., Adrion, C., Teufel, J., Bösch, S., Claassen, J., Giordano, I., Hengel, H., Jacobi, H., Klockgether, T., Klopstock, T., et al. (2017). Effects of acetyl-DL-leucine on cerebellar ataxia (ALCAT trial): study protocol for a multicenter, multinational, randomized, doubleblind, placebo-controlled, crossover phase III trial. BMC Neurol. 17, 7.

Ferber-Viart, C., Dubreuil, C., and Vidal, P.P. (2009). Effects of acetyl-DL-leucine in vestibular patients: a clinical study following neurotomy and labyrinthectomy. Audiol. Neurootol. 14, 17–25.

Fields, T., Patterson, M., Bremova, T., Belcher, G., Billington, I., Churchill, G.C., Davis, W., Evans, W., Flint, S., Galione, A., et al. (2020). A master protocol to investigate a novel therapy acetyl-L-leucine for three ultra-rare neurodegenerative diseases: Niemann-Pick type C, the GM2 Gangliosidoses, and Ataxia Telangiectasia. MedRxiv 2020.03.11.20034256.

Gleeson, M.P., Hersey, A., Montanari, D., and Overington, J. (2011). Probing the links between in vitro potency, ADMET and physicochemical parameters. Nat. Rev. Drug Discov. 10, 197–208.

Günther, L., Beck, R., Xiong, G., Potschka, H., Jahn, K., Bartenstein, P., Brandt, T., Dutia, M., Dieterich, M., Strupp, M., et al. (2015). N-acetyl-L-leucine accelerates vestibular compensation after unilateral labyrinthectomy by action in the cerebellum and thalamus. PloS One 10, e0120891.

Hall, G.G. (1991). The Lennard-Jones paper of 1929 and the foundations of Molecular Orbital Theory. Adv. Quantum Chem. 22, 1–6.

Im, H.A., Meyer, P.D., and Stegink, L.D. (1985). N-acetyl-L-tyrosine as a tyrosine source during total parenteral nutrition in adult rats. Pediatr. Res. 19, 514–518.

International Transporter Consortium, Giacomini, K.M., Huang, S.-M., Tweedie, D.J., Benet, L.Z., Brouwer, K.L.R., Chu, X., Dahlin, A., Evers, R., Fischer, V., et al. (2010). Membrane transporters in drug development. Nat. Rev. Drug Discov. 9, 215–236.

Jørgensen, K.E., Kragh-Hansen, U., and Sheikh, M.I. (1990). Transport of leucine, isoleucine and valine by luminal membrane vesicles from rabbit proximal tubule. J. Physiol. 422, 41–54.

Kaler, G., Truong, D.M., Khandelwal, A., Nagle, M., Eraly, S.A., Swaan, P.W., and Nigam, S.K. (2007). Structural variation governs substrate specificity for organic anion transporter (OAT) homologs. Potential remote sensing by OAT family members. J. Biol. Chem. 282, 23841–23853.

Kalla, R., and Strupp, M. (2019). Aminopyridines and acetyl-DL-leucine: new therapies in cerebellar disorders. Curr. Neuropharmacol. 17, 7–13.

Kanai, Y., Segawa, H., Miyamoto, K. i, Uchino, H., Takeda, E., and Endou, H. (1998). Expression cloning and characterization of a transporter for large neutral amino acids activated by the heavy chain of 4F2 antigen (CD98). J. Biol. Chem. 273, 23629–23632.

Kaya, E., Smith, D.A., Smith, C., Morris, L., Bremova-Ertl, T., Cortina-Borja, M., Fineran, P., Morten, K.J., Poulton, J., Boland, B., et al. (2020). Acetyl-leucine slows disease progression in lysosomal storage disorders. Brain Commun. fcaa148, https://doi.org/10.1093/braincomms/fcaa148.

Kaya, E., Smith, D.A., Smith, C., Boland, B., Strupp, M., and Platt, F.M. (2020). Beneficial effects of acetyl-DL-leucine (ADLL) in a mouse model of Sandhoff disease. J. Clin. Med. 9, 1050.

Keogh, J.P. (2012). Membrane transporters in drug development. Adv. Pharmacol. San Diego Calif 63, 1–42.

Krehbiel, C.R., and Matthews, J.C. (2003). Absorption of Amino Acids and Peptides. In Amino Acids in Animal Nutrition, J.P.F. D’Mello, ed. (CABI Publishing), pp. 41–70.

Leeson, P.D., and Springthorpe, B. (2007). The influence of drug-like concepts on decisionmaking in medicinal chemistry. Nat. Rev. Drug Discov. 6, 881–890.

Levy, G. (1980). Clinical pharmacokinetics of salicylates. Br. J. Clin. Pharmacol. 10, 285S–290S.

Levy, S., Kafri, M., Carmi, M., and Barkai, N. (2011). The competitive advantage of a dual-transporter system. Science 334, 1408–1412.

Lewis, G.N. (1916). The Atom and the Molecule. J Am Chem Soc 38, 762–785.

Lipinski, C.A. (2000). Drug-like properties and the causes of poor solubility and poor permeability. J. Pharmacol. Toxicol. Methods 44, 235–249.

Missner, A., and Pohl, P. (2009). 110 years of the Meyer-Overton rule: predicting membrane permeability of gases and other small compounds. Chemphyschem Eur. J. Chem. Phys. Phys. Chem. 10, 1405–1414.

Nagamori, S., Wiriyasermkul, P., Okuda, S., Kojima, N., Hari, Y., Kiyonaka, S., Mori, Y., Tominaga, H., Ohgaki, R., and Kanai, Y. (2016). Structure-activity relations of leucine derivatives reveal critical moieties for cellular uptake and activation of mTORC1-mediated signaling. Amino Acids 48, 1045–1058.

Neuzil, E., Ravaine, S., and Cousse, H. (2002). La N-acétyl-DL-leucine, médicament symptomatique de vertigineux. Bull Soc Pharm Bordx. Bull Soc Pharm Bordx. 141, 15–38.

Pade, V., and Stavchansky, S. (1998). Link between drug absorption solubility and permeability measurements in Caco-2 cells. J. Pharm. Sci. 87, 1604–1607.

Pauling, L.C. (1977). The theory of resonance in chemistry. Proc. R. Soc. Lond. Math. Phys. Sci. 356, 433–441.

Pedroso, J.A.B., Zampieri, T.T., and Donato, J. (2015). Reviewing the effects of L-leucine supplementation in the regulation of food intake, Energy balance, and clucose homeostasis. Nutrients 7, 3914–3937.

Platt, F., and Strupp, M. (2016). An anecdotal report by an Oxford basic neuroscientist: effects of acetyl-DL-leucine on cognitive function and mobility in the elderly. J. Neurol. 263, 1239–1240.

Schniepp, R., Strupp, M., Wuehr, M., Jahn, K., Dieterich, M., Brandt, T., and Feil, K. (2016). Acetyl-DL-leucine improves gait variability in patients with cerebellar ataxia-a case series. Cerebellum Ataxias 3, 8.

Schoser, B., Schnautzer, F., Bremova, T., and Strupp, M. (2019). Treatment of restless legs syndrome with acetyl-DL-leucine – accidental findings and a small case series. Eur. J. Neurol. 6, 694: EPO2251.

Strupp, M., Teufel, J., Habs, M., Feuerecker, R., Muth, C., van de Warrenburg, B.P., Klopstock, T., and Feil, K. (2013). Effects of acetyl-DL-leucine in patients with cerebellar ataxia: a case series. J. Neurol. 260, 2556–2561.

Strupp, M., Bayer, O., Feil, K., and Straube, A. (2019). Prophylactic treatment of migraine with and without aura with acetyl-DL-leucine: a case series. J. Neurol. 266, 525–529.

Truong, D.M., Kaler, G., Khandelwal, A., Swaan, P.W., and Nigam, S.K. (2008). Multi-level analysis of organic anion transporters 1, 3, and 6 reveals major differences in structural determinants of antiviral discrimination. J. Biol. Chem. 283, 8654–8663.

VanWert, A.L., Gionfriddo, M.R., and Sweet, D.H. (2010). Organic anion transporters: discovery, pharmacology, regulation and roles in pathophysiology. Biopharm. Drug Dispos. 31, 1–71.

Vibert, N., and Vidal, P.P. (2001). In vitro effects of acetyl-DL-leucine (Tanganil) on central vestibular neurons and vestibulo-ocular networks of the guinea-pig. Eur. J. Neurosci. 13, 735–748.

Waring, M.J. (2009). Defining optimum lipophilicity and molecular weight ranges for drug candidates-Molecular weight dependent lower logD limits based on permeability. Bioorg. Med. Chem. Lett. 19, 2844–2851.

van de Waterbeemd, H., Camenisch, G., Folkers, G., Chretien, J.R., and Raevsky, O.A. (1998). Estimation of blood-brain barrier crossing of drugs using molecular size and shape, and H-bonding descriptors. J. Drug Target. 6, 151–165.

Weimer, M., Jiang, X., Ponta, O., Stanzel, S., Freyberger, A., and Kopp-Schneider, A. (2012). The impact of data transformations on concentration-response modeling. Toxicol. Lett. 213, 292–298.

